# A virus-induced circular RNA maintains latent infection of Kaposi sarcoma herpesvirus

**DOI:** 10.1101/2022.07.18.500467

**Authors:** Takanobu Tagawa, Daniel Oh, Sarah Dremel, Guruswamy Mahesh, Vishal N. Koparde, Gerard Duncan, Thorkell Andresson, Joseph M. Ziegelbauer

**Affiliations:** HIV and AIDS Malignancy Branch, Center of Cancer Research, National Cancer Institute; Bethesda; Maryland, USA; CCR Collaborative Bioinformatics Resource, Center for Cancer Research, National Cancer Institute, National Institutes of Health, Bethesda, Maryland, USA; Advanced Biomedical Computational Sciences, Frederick National Laboratory for Cancer Research, Leidos Biomedical Research, Inc., Frederick, Maryland, USA; Protein Characterization Laboratory, Cancer Research Technology Program, Frederick National Laboratory for Cancer Research, Leidos Biomedical Research, Inc., Frederick, Maryland, USA

## Abstract

Non-coding RNAs (ncRNAs) play important roles in host-pathogen interactions; oncogenic viruses like Kaposi sarcoma herpesvirus (KSHV) employ ncRNAs to establish a latent reservoir and persist for the life of the host. We previously reported that KSHV infection alters a novel class of RNA, circular RNAs (circRNAs). CircRNAs are alternative splicing isoforms and regulate gene expression but, their importance in infection is largely unknown. Here, we showed that a human circRNA, hsa_circ_0001400, is induced by various pathogenic viruses, namely KSHV, Epstein-Barr virus, and human cytomegalovirus. The induction of circRNAs including circ_0001400 by KSHV is co-transcriptionally regulated, likely at splicing. Consistently, screening for circ_0001400-interacting proteins identified a splicing factor, PNISR. Functional studies using infected primary endothelial cells revealed that circ_0001400 inhibits KSHV lytic transcription and virus production. Simultaneously, the circRNA promoted cell cycle, inhibited apoptosis, and induced immune genes. RNA-pulldown assays identified transcripts interacting with circ_0001400, including *TTI1*, which is a component of the pro-growth mTOR complexes. We thus identified a circRNA that is pro-growth and anti-lytic replication. These results support a model in which KSHV induces circ_0001400 expression to maintain latency. Since circ_0001400 is induced by multiple viruses, this novel viral strategy may be widely employed by other viruses.

**AUTHOR SUMMARY:** Circular RNAs (circRNAs) are single-stranded, closed-circular RNAs. They coincide with linear mRNAs as alternative splice isoforms. CircRNAs binds to other RNAs or proteins to regulate their abundance or functions. We previously showed that specific human circRNAs are differentials regulated upon infection with an oncogenic DNA virus, Kaposi’s sarcoma herpesvirus (KSHV). Functions of such circRNAs for virus infections were largely elusive. Here, we identified that one of circRNAs, hsa_circ_0001400 controls both viral and human gene expression so that infected human cells have less virus production but better cell growth. KSHV, like other herpesviruses, has two phases: latent and lytic cycle. The functions of circ_0001400 suggest it biases the cells to latent cycle during which, unlike lytic phase, viruses express only limited number of genes to survive and new viruses are not produced. Since oncogenic viruses like KSHV mainly stay at latent phase for the life of the host, our results suggest that the virus utilized circRNAs to switch to latent phase after the infection. Infection of other pathogenic viruses like Epstein-Barr virus and human cytomegalovirus were also found to induce circ_0001400. The circRNA may be thus controlling infection of various viruses.

## INTRODUCTION

Oncogenic DNA viruses like Kaposi’s sarcoma herpesvirus (KSHV) and Epstein-Barr virus (EBV) establish lifelong infection in humans. Around 20% of cancers are associated with oncogenic viruses, and KSHV and EBV are associated with various cancers including Kaposi’s sarcoma, primary effusion lymphoma (PEL), Burkitt’s lymphoma, and nasopharyngeal carcinoma (Shannon-Lowe and Rickinson, 2019). Though primary infections are thought to happen in B lymphocytes, cell types of associated cancers can include B cells, T cells, epithelial cells, and endothelial cells. A crucial strategy for gamma herpesviruses, KSHV and EBV, is to establish latent infection during which most of viral genes are silenced and therefore less immunogenic. During the lytic phase, in contrast, reactivated virus expresses upwards of 80 transcripts, replicate their genome, and produce infectious progeny. Lytic replication is critical for viral spread; this, however, is at the expense of the host cells. For long-term survival of the virus and infected cells as in the case of KSHV or EBV, maintaining latency is the key strategy.

Non-coding RNAs (ncRNAs) are increasingly deemed important to control infection. For example, Hepatitis C virus depends on a human microRNA (miRNA), hsa-miR-122, for viral RNA accumulation (Sarnow and Sagan, 2015) and KSHV’s long non-coding RNA, PAN, is required for virus productions (Rossetto and Pari, 2014). Viruses also exploit miRNAs to maintain latent infection; hsa-miR-155 is induced by EBV and mimicked by KSHV infection to support latent infection (Lu et al., 2008; Skalsky et al., 2007; Yin et al., 2008). Circular RNAs (circRNAs) are novel in the interactions between host and viruses. CircRNAs are alternative splice isoforms produced from back-splicing. They are single-stranded closed circular RNAs with the ability to interact with other RNA species or proteins to regulate their functions (Kristensen et al., 2019). Some of the first reports of circRNAs demonstrated inhibition of miRNA activity by a circRNA (Hansen et al., 2013; Memczak et al., 2013). The importance of circRNAs is particularly evident in cancers. Numbers of human circRNAs have been characterized to be oncogenic (e.g. circPVT1, CDR1as) or anti-oncogenic (e.g. circsMARCA5 or circSHPRH)(Kristensen et al., 2019).

The contributions of human circRNAs during viral infection are largely unknown. We previously found that KSHV *de novo* infection differentially regulates dozens of human circRNAs and that the virus encodes its own circRNAs (Tagawa et al., 2021, 2018; Toptan et al., 2018; Ungerleider et al., 2019). One of them, hsa_circ_0001400 was induced upon infection with KSHV and, in a renal carcinoma cell line, reduced KSHV gene expression. The importance of human circRNAs in viral infection is still elusive, particularly in clinically relevant settings such as primary endothelial cells: Does hsa_circ_0001400 control latent/lytic infection? Is host cell physiology affected by circRNAs? What is the mechanism of gene regulation by the circRNA? How common is the induction of circ_0001400 by viruses other than KSHV?

Here, we performed high throughput circRNA profiling in different herpesviruses and cell types with a particular focus on *de novo* infection. Functions of circ_0001400 were assessed in human primary endothelial cells regarding viral transcription, virus production, host gene regulation, phenotype of infected cells, and immune responses. To identify RNAs and proteins that interact with circ_0001400, we performed circRNA-RNA pulldown assays as well as proximity labeling-based circRNA-protein pulldown experiments. All in all, our results encompass that circ_0001400 functions to maintain KSHV latent infection.

## RESULTS

### Infection of multiple herpesviruses induces the human circRNA, hsa_circ_0001400

A human circRNA, hsa_circ_0001400 (circ_0001400, circRELL1), encoded from the *RELL1* gene locus, increases upon *de novo* KSHV infection in epithelial cell lines as well as human primary endothelial cells including human umbilical vein endothelial cells (HUVECs) (Tagawa et al., 2018) and lymphatic endothelial cells (LECs) (Fig. 1A). Circ_0001400 was one of most abundant circRNAs in HUVECs as well (Fig. 1B). We found that circ_0001400 is induced upon infection with various human herpesviruses. The circRNA was more abundant in the Epstein-Barr virus-positive Burkitt’s lymphoma cell line Akata(+) than in an episome-negative clone Akata(-) (Fig. 1A). Similar to KSHV infection, human cytomegalovirus infection increased circ_0001400 levels in the MRC5 fibroblast cell line (Fig. 1A)(Oberstein and Shenk, 2017). Thus, increased levels of circ_0001400 were found after infection with multiple pathogenic human herpesviruses.

**Fig. 1.**
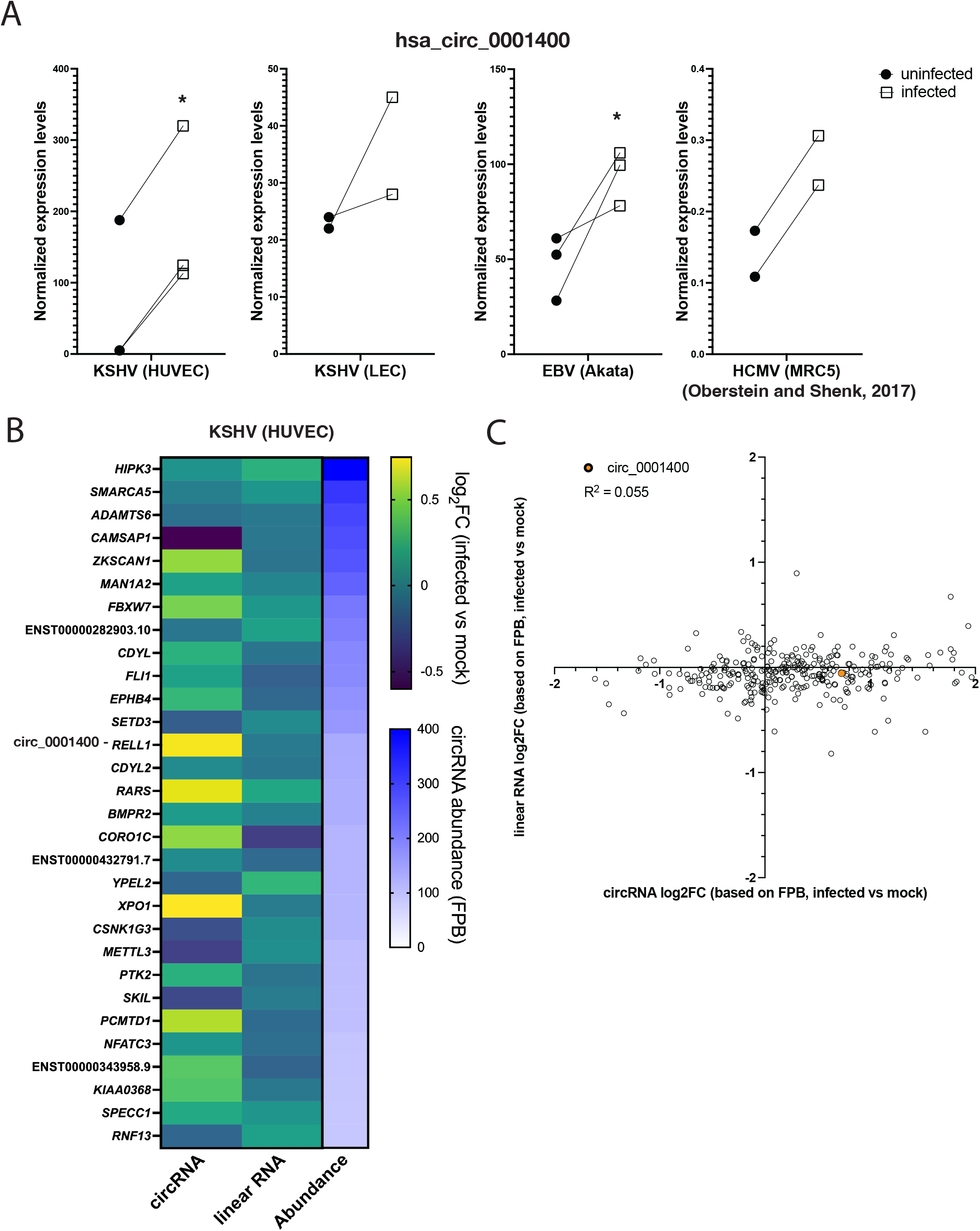
Herpesviruses induce circ_001400 expression. (A) hsa_circ_0001400 transcript levels are shown in KSHV-infected HUVECs (microarray), LECs (RNA-Seq), EBV-positive Burkitt’s lymphoma line Akata (microarray), and HCMV-infected fibroblast (RNA-Seq)(Oberstein and Shenk, 2017). n=2-3. Significances were computed with limma. *:p-value < 0.05. (B) Fold-changes of top 30 most abundant human circRNAs in KSHV-infected HUVECs. Transcript levels of circRNAs and corresponding linear mRNAs were quantitated from total RNA-Seq with circExplorer3 (Ma et al., 2019) as FPBs (fragments per billion mapped base). Fold-changes of circRNAs and linear RNAs (infected vs mock) are calculated (n=3) and mean values are shown. Genes are sorted by circRNA FPBs (Abundance) and host genes of both circRNA and linear RNAs are shown on the left. FPBs and ratios are available in Table S1. (C) Log_2_ fold-changes of all detected human circRNAs/counterpart linear mRNAs (KSHV-infected vs mock HUVECs) are shown as a scatter plot. Data is same as in Fig. 1B and Table S1. Pearson correlation value was calculated. n=3 and mean values are shown.

How viral infection regulates circRNA transcript levels is elusive. The amount of human circRNA may be regulated by altering transcription rates but also by modulating co-transcriptional steps, particularly back-splicing, which is known to be regulated by RBPs (Kristensen et al., 2019). To determine how KSHV regulates circRNA production, we re-analyzed our previous datasets to quantitate circRNAs and counterpart linear mRNAs in KSHV-infected endothelial cells (Fig. 1B-C) (Serquiña et al., 2021). While circ_0001400 is increased by infection, the mRNA from same gene locus, *RELL1* was not differentially regulated along with other circRNAs like circZKSCAN1, circRARS, and circXPO1(Fig. 1B). This lack of the correlation between fold-changes of circRNAs and linear RNAs are, in fact, global with a Pearson correlation coefficient of 0.055 (Fig. 1C, Table S1). These results show that KSHV infection changes the maturation, localization, or stabilities of circRNAs rather than altering transcription rates of circRNA-encoding genes.

### hsa_circ_0001400 are regulated co-transcriptionally and interact with a splicing factor

To further investigate the nature of circRNA regulation, we performed 4SU RNA-Seq. Conventional total RNA-Seq quantitates steady-state level of transcripts while 4SU-Seq only measures newly transcribed RNAs (Fig. 2A). This technique enables us to discern whether KSHV globally alters host circRNA abundance by influencing co-transcriptional (e.g. splicing) or post-transcriptional (e.g. RNA decay) processes. Upon reactivation of KSHV-positive iSLK-BAC16 cells, the circRNA:linear RNA ratio of circ_0001400 increased more in 4SU-labeled (nascent) RNAs than total RNAs (Fig. 2B) indicating co-transcriptional increase by KSHV. Most human circRNAs were differentially regulated in a co-transcriptional manner (Fig. 2B). This is consistent with the observation in HUVECs where changes of circRNA and linearRNA upon *de novo* infection did not correlate (Fig. 1B-C). During reactivation, 75% of circRNAs were classified to be co-transcriptionally regulated including circ_0001400 (Fig. 2B).

**Fig. 2.**
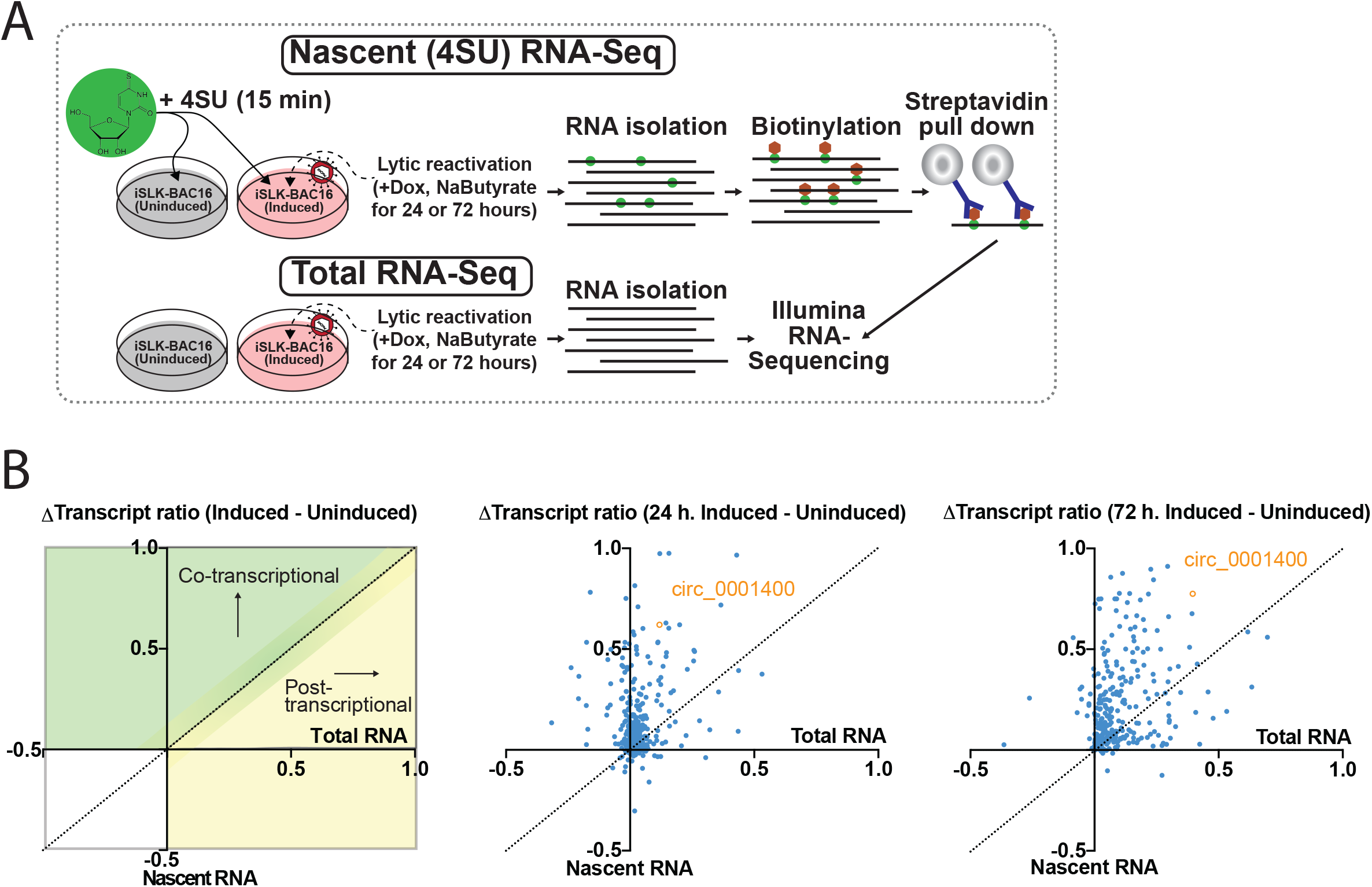
circ_001400 is co-transcriptionally regulated and interacts with a splicing factor. (A) Infographic for sequencing techniques performed. (B) Total and 4SU RNA-Seq delta transcript ratio (circular reads/linear reads) for unique human circRNA (1 or 3 d). Positive Δtranscript ratios (Induced – Uninduced) indicate a shift after lytic reactivation in transcript abundance, favoring the circular transcript. circ_001400 is highlighted in orange. n=2 and mean values are shown.

The back-splicing step of circRNA maturation is often controlled by RNA-binding proteins (Kristensen et al., 2019). We utilized a CRISPR-assisted RNA–protein interaction detection method, CARPID (Fig. 3A)(Yi et al., 2020) to identify proteins interacting with circ_0001400. CRISPR–Cas13 based targeting of circRNAs is specific and efficient (Li et al., 2021). Guide RNAs targeted against circ_0001400 could indeed knockdown the circRNA whereas the counterpart mRNA RELL1 is minimally affected (Fig. 3B). Since CARPID depends on proximity labeling, we can identify proteins that are RBPs directly bound to circ_0001400 as well as indirectly interacting proteins with mass spectrometry. We identified various proteins including RBPs (Fig. 3C and Table S2) and confirmed the interaction between circ_0001400 and PNISR, but not EIF3K, with RNA immunoprecipitation assays (Fig. 3D). Since PNISR is a splicing factor and circRNAs are formed with back-splicing, PNISR may play a role in regulating specific circRNAs such as circ_0001400. The 4SU RNA-Seq analysis and identification of a circRNA-interacting splice factor thus suggesting that KSHV regulates human circRNAs mostly co-transcriptional manner, including circ_0001400.

**Fig. 3.**
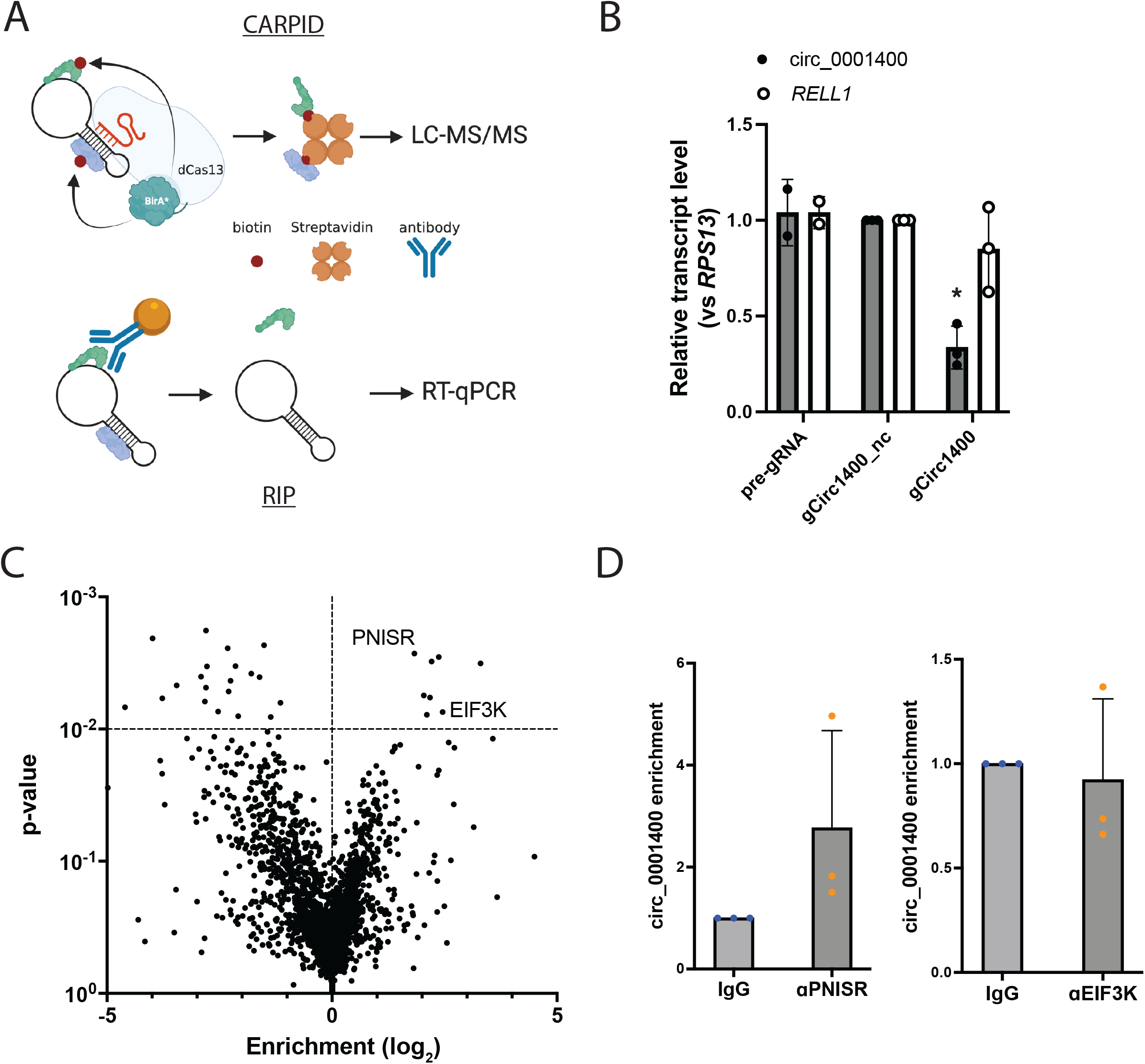
Protein interactome of circ_0001400. (A) Diagram of CRISPR-assisted RNA–protein interaction detection method (CARPID) and RNA-immunoprecipitation (RIP) method. CARPID biotinylate proteins that are in the proximity of circ_001400 to enrich and determine their identities by liquid chromatography/tandem mass spectrometry (LC-MS/MS). Identified proteins are listed in Table S2. Candidate proteins are immunoprecipitated by specific antibodies (RIP) and circ_001400 is confirmed to be co-precipitated by RT-qPCR. (B) Confirmation of specificity and efficacy of guide RNAs. 293T cells were transfected with expression plasmids harboring a pre-gRNA, a negative control gRNA, or a circ_0001400-specific gRNA and CasRx-expression plasmid for 48 hours. RT-qPCR was performed to measure transcript levels and normalized to the negative control gRNA result (n=2-3). Significances were calculated with paired-t tests to the control when n=3. *:p-value < 0.05. (C) A volcano plot of proteins that were identified by mass spectrometry (n=3). Enrichment and significances were calculated by RankProd package. Pre-gRNA (empty) vector and gCirc1400_nc vector were used as negative controls. (D) RIP results of candidate proteins. circ_0001400 was ectopically expressed in 293T cells for 24 hours and RIPs were performed against PNISR and EIF3K. circ_001400 transcript levels after RIP are normalized to the isotype IgG antibody condition. n=3 and significances were calculated with paired-t tests. *:p-value < 0.05.

### KSHV lytic gene expression and virus production are inhibited by circ_0001400

Effects of circ_0001400 early after KSHV infection were assessed using lentiviral transduction and RNAi-mediated knockdown of the circRNA (Fig. 4, Fig. S1, Table S3). The lentivirus vector expresses a cassette that harbors circ_0001400 flanked with ZKSCAN1 introns, which facilitates back-splicing and circularization of RNAs (Liang and Wilusz, 2014). In accordance with our observation with infected SLK cells (Tagawa et al., 2018), a renal carcinoma cell line, ectopic expression of circ_0001400 (Fig. S1A) reduced KSHV gene expression in HUVECs (Fig. 4A-B). Viral transcriptome analysis after ectopic expression of circ_0001400 in HUVECs showed suppression of gene expression for most of the KSHV genes (Fig. 4A). The siRNA specifically targets circ_0001400 back-splice junctions, which is unique to circRNAs. The siRNA was proven to be specific to the circRNA and does not affect its counterpart mRNA (Tagawa et al., 2018). Consistently, knock-down of circ_0001400 with a circRNA-specific siRNA (Fig. S1A) resulted in an increase of viral transcripts (Fig, S1C and Table S3). We observed a difference between lytic and latent genes in suppression: while lytic genes are suppressed by log_2_FC of -1.01±0.25, latent genes (K1, K2, K12, K13, LANA, and ORF72)(Arias et al., 2014) were reduced only by log_2_FC of -0.20±0.14 (Fig. S1C and Table S3). Importantly, manipulation of circ_0001400 expression levels did not affect infectivity, suggesting viral gene expression is regulated at a post-entry step (Fig. 4C and S1D).

**Fig. 4.**
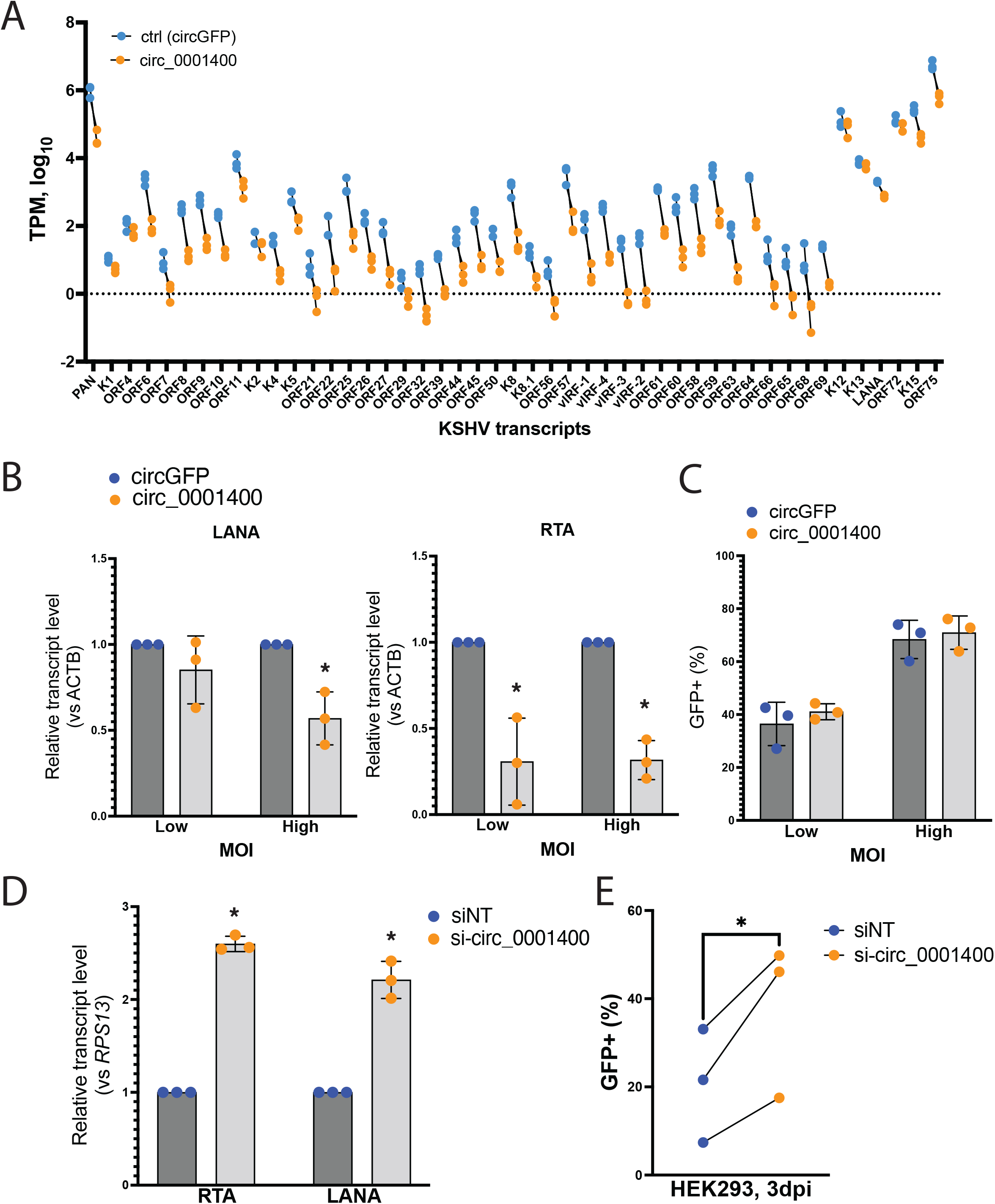
circ_001400 inhibits viral gene expression and virion production. (A) Viral transcriptome analysis after circ_0001400 ectopic expression in KSHV-infected HUVECs (3 days post infection). n=3. TPMs (Transcripts Per Kilobase Million) were calculated and only KSHV genes with significant read counts are shown. (B) Viral transcript levels after circ_0001400 ectopic expression in KSHV-infected HUVECs. MOI of 0.5 (low) and 1.0 (high) are used and RNAs were collected 3 days post infection (n=3). (C) Populations of GFP-positive, KSHV infected among lentivirus-infected HUVECs are shown. Cells were co-infected with KSHV and lentivirus (circGFP control or circ_0001400) for 3 days (n=3). GFP signal from circGFP is significantly lower than KSHV BAC16-originated GFP and does not affect the analysis. (D) Viral transcript levels after circ_0001400 ectopic expression in KSHV-infected LECs. MOI of 1 was used for infection and total RNAs were collected 3 days post infection (n=3). (E) Virus production from circ_0001400-depleted KSHV-infected LECs. Conditioned media of KSHV-infected LECs (D) was collected and used for infection of HEK-293 cells. KSHV-positive, GFP-positive population among HEK-293 cells were measured 3 days post infection (n=3). Significances were calculated with paired-t tests. *:p-value < 0.05.

Since lytic genes are more sensitive to circ_0001400, we evaluated the effect of changing circ_0001400 levels in virus production assays. Since HUVECs are latent infection model, we used LECs, which spontaneously enter lytic phase after *de novo* infection (Cheng et al., 2011). Expression of circ_0001400 was depleted by siRNAs (Fig. S1B) and conditioned media was collected 3 days post infection. To quantitate functional virions secreted from infected LECs, the conditioned media was used to infect to HEK-293 cells and KSHV-infected populations were measured using the GFP reporter expressed by the virus. Depletion of the circRNA lead to a significant increase in viral gene expression in LECs as in HUVEC infections (Fig. 4D) as well as newly produced virions (Fig. 4E). Taken together, these results suggest that circ_0001400 can suppress viral lytic infection in primary endothelial cells.

### Circ_0001400 promotes cell cycle, inhibits apoptosis, and induces genes involved in immunity

Human circRNAs can regulate expression of multiple genes to control cell physiology (Kristensen et al., 2019). To assess host genes affected by circ_0001400, we performed transcriptome analysis of circRNA-manipulated HUVECs (Fig. 5). We observed that 184 genes were consistently and differentially expressed genes (DEGs) by gain- and loss-of-function manipulations of circ_0001400 (Fig. 5A and Table S4). DEGs include genes involved in cell growth, immune responses, and crucial miRNA components, *AGO1* and *AGO2*, suggesting that circ_0001400 may behave as a regulator of the miRNA machinery (Fig. 5A). Gene enrichment analysis showed pro-cell growth tendency of circ_0001400 with upregulation of PI3K/AKT signaling and downregulation of PTEN signaling and GADD45 signaling (Fig. 5B). Since these regulations suggests changes in cell cycle and apoptosis, we evaluated these phenotypes. We found circ_0001400-depletion indeed caused G2 cell cycle arrest and an increase of apoptosis in endothelial cells (Fig. 5C-D), confirming the effects of this circRNA on pro-growth outcomes.

**Fig. 5.**
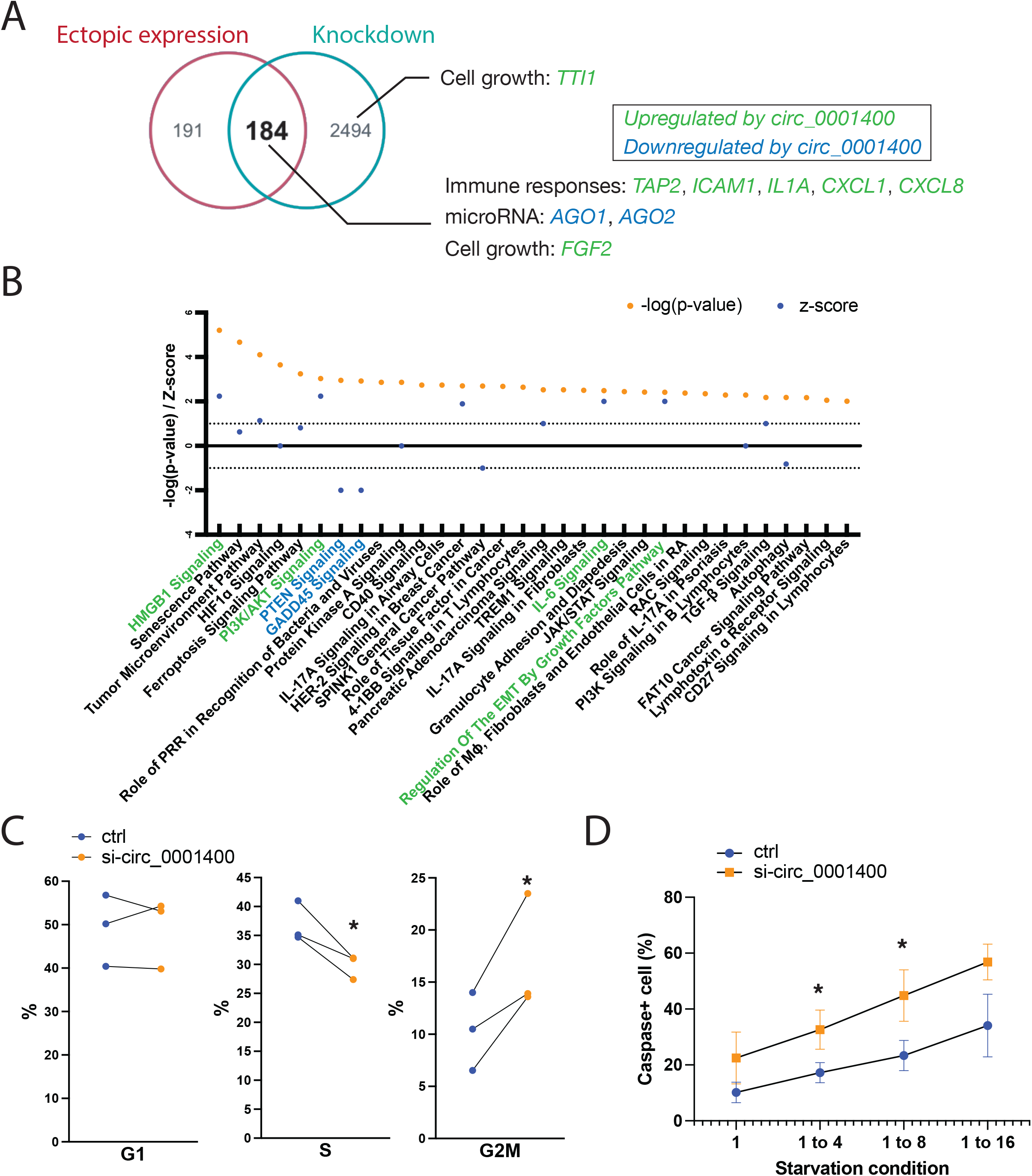
circ_001400 promotes cell growth and induces immune reaction genes. (A) Venn diagram is showing consistently differentially regulated genes (DEGs) by circ_0001400 in HUVECs. Cells were treated with siRNA targeted against circ_0001400 (n=3) or infected with circ_0001400-expressing lentivirus (n=3) followed by KSHV infection. DEGs are defined as having adjusted p-value <0.05 as calculated by voom (limma package). On 3 days post infection, the total RNA was extracted for RNA-Seq. Select up-regulated or down-regulated genes by circ_0001400 are shown in green and blue, respectively. The list of DEGs is available in Table S4. (B) Pathway enrichment analysis of DEGs (A) by Ingenuity Pathway Analysis (Qiagen). Significance of pathway enrichment is shown as p-values. Z-scores describe the predicted activation or inhibition of pathways. Pathways are color-coded when |Z| > 1.96. (C) Cell cycle analysis of circ_0001400-depleted KSHV-infected cells. Infected cells were incubated with EdU at 3 days post infection for 4 hours and cell cycles were quantitated by flow cytometry (n=3). (D) Apoptosis assay of circ_0001400-depleted KSHV-infected cells. Cells were siRNA-treated and infected. The next day, the media was replaced with diluted media to the ratio as described. At 3 days post infection, Caspase3/7 substrates that emit signal upon cleavage were added to measure apoptosis rates by high-content imaging (n=3). Significances were calculated with paired-t tests. *:p-value < 0.05.

Simultaneously, activation of immunity-related pathways including HMGB1 signaling and IL-6 signaling, was correlated with circ1400 expression. Notably, DEGs include key genes involved in antigen presentation like *TAP2* and *ICAM1* as well as various inflammatory cytokines and chemokines (Fig. 5A). ICAM1 is one of the co-stimulatory molecules, which are necessary to activate antiviral T-cells. Depletion of circ_0001400 resulted in reduced transcript levels (Fig. 6A) as well as cell surface expression levels of co-stimulatory molecules for antigen presentation including ICAM1 (Fig. 6B).

**Fig. 6.**
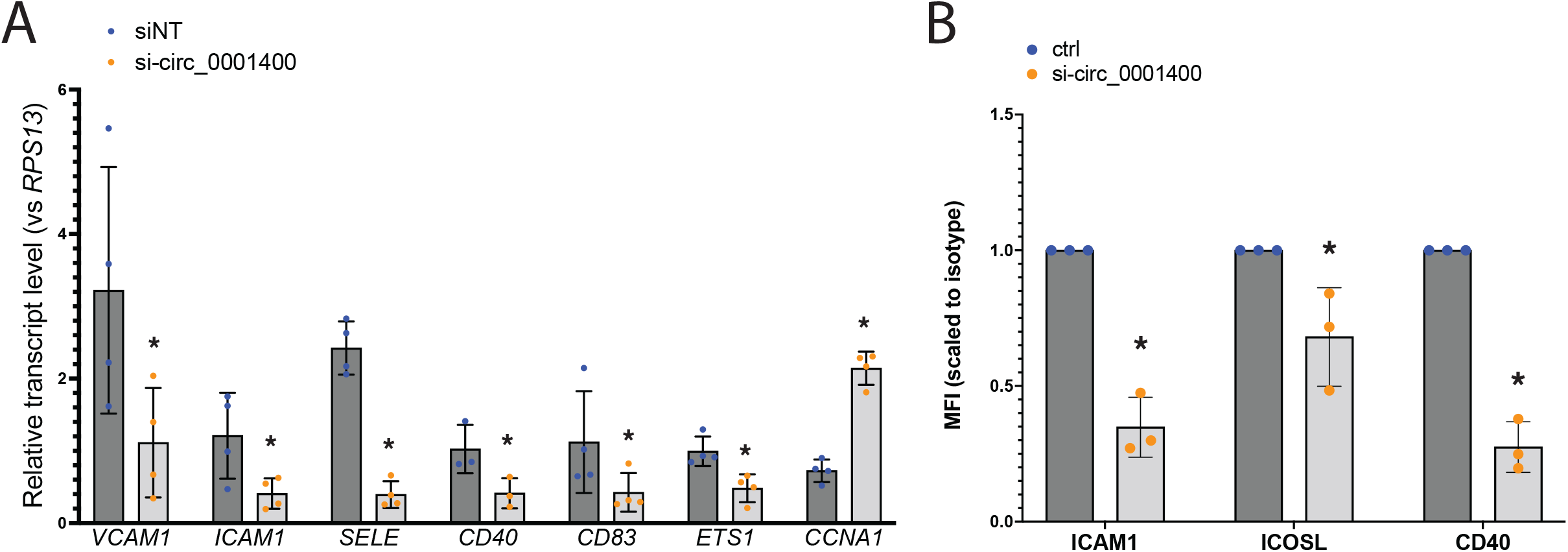
circ_001400 activates immune gene expression and increases co-stimulatory molecules on cell surface. (A) Human transcript levels after circ_0001400 ectopic expression in KSHV-infected HUVECs (MOI 0.5). Total RNAs were extracted 3 days post infection for RT-qPCR (n=3). Significances were calculated with paired-t tests. *:p-value < 0.05. (B) Immune staining of co-stimulatory molecules on circ_0001400-depleted KSHV-infected cells. At 3 dpi, cells were stained with antibodies specific to co-stimulators and isotypes (n=3). Mean fluorescence intensity (MFI)for each antibody was scaled to the signal from isotypes and shown. Significances were calculated with paired-t tests. *:p-value < 0.05.

### Circ_0001400 interacts with mTOR component and a splicing factor

RNA-RNA interactions are one mechanism circRNAs use to alter gene regulation (Kristensen et al., 2019). We searched for RNAs associated with circ_0001400 using RNA-pulldown assays in 293T cells. A biotinylated DNA oligo that is complementary to the back-splice junction of circ_0001400 was used to pull-down circ1400 and interacting RNAs were determined by deep-sequencing (Fig. 7A). To increase the sensitivity, cells were treated with psoralen followed by long-wave UV radiation, which covalently crosslinks double-stranded RNAs. Extracted RNAs were exposed to short-wavelength UV to reverse the cross-linking. The top candidate of interacting RNAs were shown in Table S5.

**Fig. 7.**
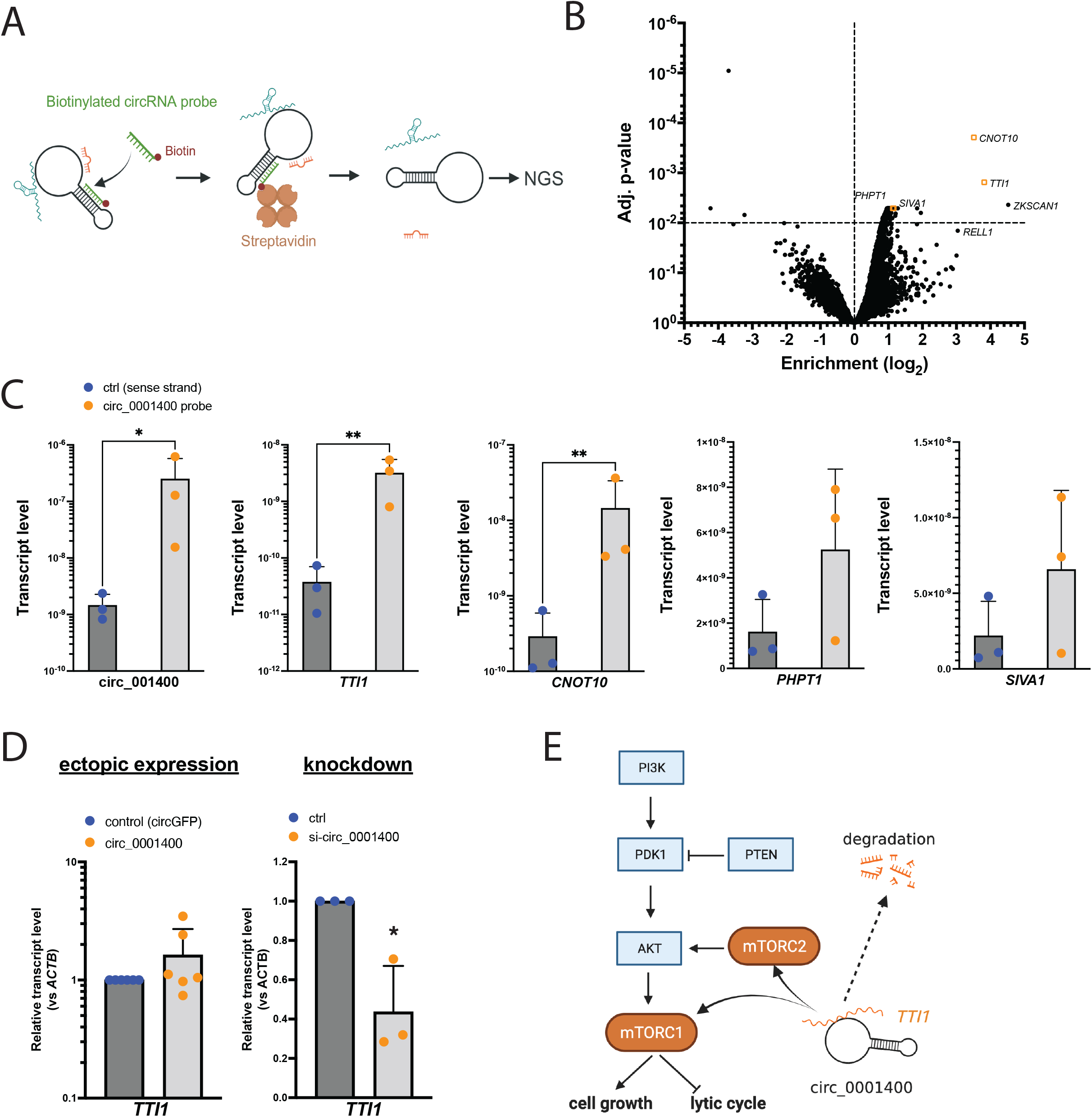
circ_001400 interacts with mTOR-component *TTI1* and sustains its transcript level. (A) Diagram of the RNA-pulldown assay is shown. Total RNA was incubated with biotinylated DNA oligos that is specific to circ_0001400 and circRNA-RNA complexes are purified using streptavidin beads. Next generation sequencing (NGS) was performed to determine their identities. (B) A volcano plot of transcripts identified by the circ_0001400 RNA pulldown assay in 293T cells. Mapped reads were median-scaled and enrichments were calculated by voom (limma package). *ZKSCAN1* and *RELL1* are partially included in the circ_0001400 expression plasmid vectors and thus serve as positive control of pulldown. Transcripts that were confirmed in Fig. 6C are depicted as orange rectangles. n=3. (C) RT-qPCR of enriched transcripts after RNA-pulldown. circ_0001400 was ectopically expressed in 293T cells for 24 hours and RNA-pulldowns were performed (n=3). Enrichment of circ_0001400 and candidate mRNAs (listed in Table S5) are shown. (D) *TTI1* transcript levels after manipulation of circ_0001400 transcript levels. circ_0001400 was ectopically expressed (n=6) or depleted by siRNAs (n=3) in 293T cells for 48 hours. Total RNAs were extracted followed by RT-qPCR. (E) Proposed model of circ_0001400’s regulation of PI3K/AKT/mTOR pathway. Significances were calculated with paired-t tests. *:p-value < 0.05.

*TTI1* and *CNOT10* mRNAs showed the highest enrichment as well as adjusted p values (Fig. 7B). TTI1 is a component of mTOR (mammalian target of the rapamycin) complex and important in forming mTOR complexes (Kaizuka et al., 2010; Pal et al., 2021; Takai et al., 2010), which is known to promote cell growth and inhibit lytic cycle of KSHV (Purushothaman et al., 2015). CNOT10 is a component of the carbon catabolite repression 4 (CCR4)–negative on TATA-less (NOT) complex, which serves as a deadenylase and is critical for miRNA-mediated mRNA degradation (Shirai et al., 2014). The interaction between circ1400 and these RNAs was confirmed with RT-qPCR (Fig. 7C). *PHPT1* and *SIVA1* showed only a mild enrichment (Table 2); RT-qPCR showed tendency of enrichment value but not statistically significant (Fig. 7B), indicating the use of both enrichment value and p value is appropriate for finding interacting RNAs.

Since circ_0001400 activated PI3K/AKT signaling pathway and showed pro-growth, anti-lytic cycle phenotype, the regulation of *TTI1* mRNA levels by circ_0001400 was assessed (Kaizuka et al., 2010; Pal et al., 2021; Takai et al., 2010)(Purushothaman et al., 2015)(Shirai et al., 2014). Depletion, but not ectopic expression, of circ_0001400 significantly alter the transcript level of *TTI1* in 293T (Fig. 7D). The same tendency was observed in primary endothelial cells (Fig. 5A and Table S4). These results suggests that circ_0001400 helps to maintain *TTI1* mRNA levels. These results suggest that circ_0001400 interacts with *TTI1* to sustain its transcript levels (Fig. 7E). We thus identified RNAs that may account for pro-growth, inflammatory, and anti-lytic phenotypes caused by circ_0001400 in KSHV-infected cells.

## DISCUSSION

In KSHV-infected primary human endothelial cells, we found that circ_0001400 inhibited lytic gene expression and virus production while the circRNA also promoted cell cycle and suppressed apoptosis. Though the former phenotypes apparently are antiviral and the latter pro-growth feature are oncogenic and pro-viral, these phenotypes support the notion that: circ_0001400 propagates infected cells that are in latency, during which only handful of viral genes (latent genes) are expressed and viruses are not produced (Fig. 8). Switching to latent infection after infection is critical for herpesviruses like KSHV and EBV since this less immunogenic state allows viruses to evade immune surveillance and persist for the life of host. These wide range of phenotypes caused by circ_0001400 in primary endothelial cells are in stark contrast to our previous study in a renal carcinoma cell line, SLK (Tagawa et al., 2021). There, we observed the inhibition of viral gene expression, but cell growth or immune genes were not affected and thus the circRNA conferred antiviral functions. This also signifies the importance to utilize models that are closer to clinical conditions, such as primary endothelial cells and *de novo* infection system.

**Fig. 8.**
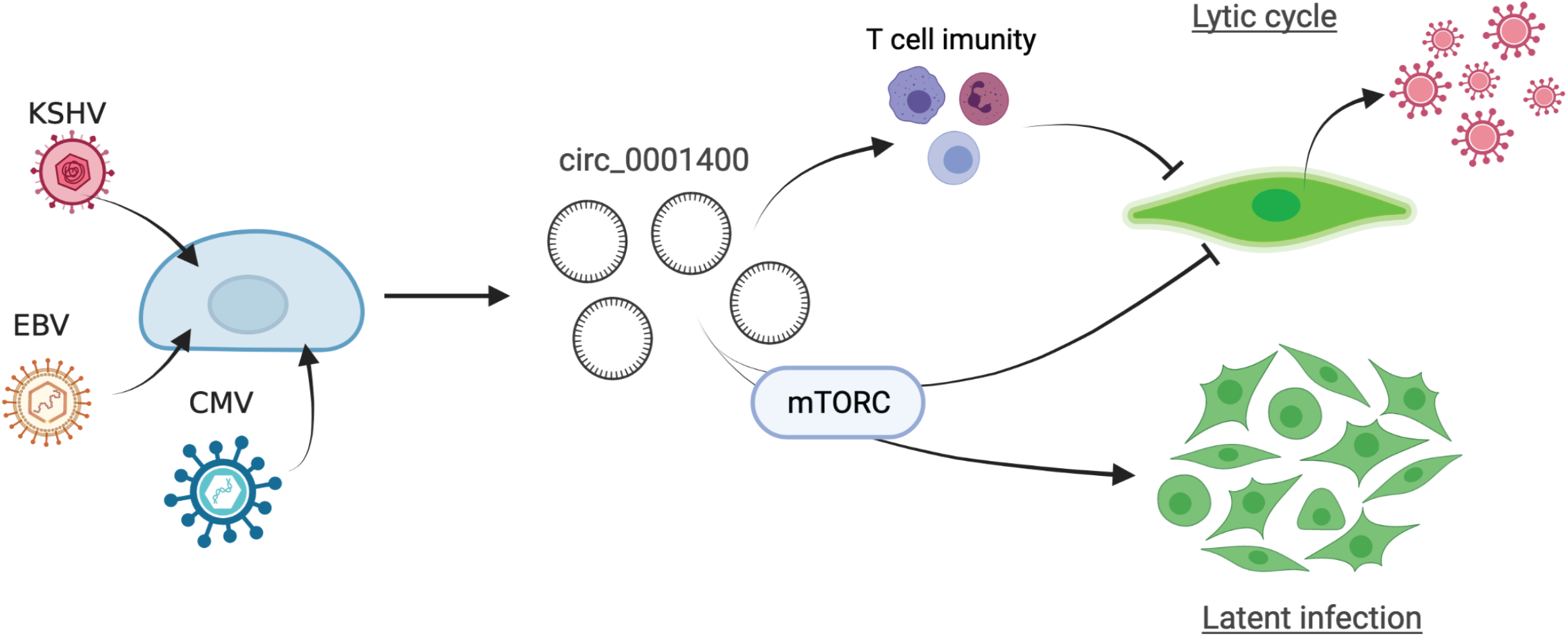
Model of hsa_circ_0001400 in virus-host interactions. hsa_circ_0001400 is induced by multiple herpesviruses. The circRNA allows activation of PI3K/AKT/mTOR pathway and promote cell growth of latently KSHV-infected cells. On the other hand, lytic infection is suppressed by inhibition of virus production and increase of co-stimulatory molecules by circ_0001400.

We identified hsa_circ_0001400 to be induced by various human herpesviruses, KSHV, EBV, and HCMV. It is also noteworthy that the increase of the circRNAs occurs in wide range of cell types from epithelial cells, endothelial cells, fibroblasts, and B lymphocytes. Since circ_0001400 is just one of dozens of infection-induced circRNAs, these observations present the following question: Are there any other circRNAs also regulated in infection with various viruses and cell types? And, if so, what will be the aggregate effect of circRNAs in host-virus interactions? Circ_0001400 alone has already showed a significant role in cell growth and latency, and the potential of circRNAs, and non-coding RNAs as extension, should be highlighted.

Stimulating the PI3K/AKT/mTOR pathway is critical for KSHV infection. The pathway is simulated by multiple viral proteins, like K1, vGPCR (ORF74), vIL-6 (K2), and ORF45. KSHV induces the PI3K pathway upon *de novo* infection to promote cell growth and reduce apoptosis (Bhatt and Damania, 2013) as well as maintaining latency (Peng et al., 2010). mTOR is also known to induce immune genes like ICAM1 via NF-kB pathway in HUVECs (Minhajuddin et al., 2005). Consistently, we observed same pro-growth phenotypes, activation of immune genes, and inhibition of lytic cycles by circ_0001400. RNA-pulldown assay showed the circRNA’s interaction with a mTOR component, *TTI1*, and circ_0001400 sustains transcript levels of TTI1 in 293T cell and HUVECs. Further, *TTI1* is one of PEL-specific oncogenic dependencies (Manzano et al., 2018) indicating its importance in KSHV-driven oncogenesis. These results suggests that circ_0001400 causes pro-latency phenotypes via maintaining PI3K/AKT/mTOR pathway.

Immune regulatory effects of circ_0001400 was suggested by transcriptomic data. Co-stimulators of antigen presentations such as ICAM1 are well documented to be suppressed by KSHV (Lee et al., 2012). On the other hand, induction of circ_0001400 upon KSHV infection increased co-stimulators including ICAM1, which may lead to higher immunogenicity and removal of infected cells by host immune surveillance system. In addition, genes coding for inflammatory cytokines and chemokines, *IL1A, CXCL1, CXCL8* (coding for IL-8) are also induced by circ_0001400. This may suggest fine-tuning of immunogenicity by host circRNAs. Though immune evasion by reducing antigen presentation is important and necessary, removal of excessively immunogenic cells i.e. lytic cells, potentially via circ_0001400, is beneficial to maintain latent infection *in vivo* by avoiding anti-viral inflammatory microenvironment at infection sites (Fig. 8).

RNA binding proteins were recently found to interact with certain circRNAs (Huang et al., 2020). Our CARPID dataset therefore can be utilized to search for such proteins interacting with circ_0001400 in future studies. Among enriched circRNA-interacting proteins, one apparent omission was the Argonaut family of proteins, core proteins of RNA-induced silencing complex (RISC). This was unexpected since some circRNA regulatory mechanisms include acting as decoys or “sponges” for miRNAs. It is possible that because circ_0001400 is relatively short at 434 nucleotides, there are no enriched binding sites of particular miRNAs. Well studied miRNA sponge circRNA CDR1as is 1.5 kb and harbors ∼70 miR-7 binding sites. *In silico* prediction did not find such concentrated miRNA binding sites of circ_0001400. Since circ_0001400 was suggested to control *AGO1, AGO2*, and *CNOT10*, this circRNAs may regulate general miRNAs machinery, rather than targeting specific miRNAs.

Another surprising observation on the circ_0001400’s interactions is, at least for *TTI1*, depletion of circ_0001400 reduced the amount of the interacting RNAs (Kristensen et al., 2019). Though circRNAs are conventionally known to suppress bound RNAs, there is an example that circRNAs maintain levels of a bound miRNA (Piwecka et al., 2017). It is unclear whether circ_0001400 binds to mRNAs directly or indirectly yet, but similar circRNA-mediated RNA maintenance may play a role for the circ_0001400’s gene regulatory mechanism.

We also explored the mechanism by which KSHV infection increased the amounts of circRNAs. Here, we show that most of differentially regulated circRNAs by infection are regulated at co-transcriptional levels, including circ_0001400, likely at back-splicing steps. We have previously shown that circ_0001400 synthesis depends on multiple RBPs like FUS and QKI (Tagawa et al., 2021). We newly identified PNISR to interact with circ_0001400. Since PNISR is a splicing factor, it is another candidate to be involved in the synthesis of circ_0001400. If PNISR and other candidates are specific to circ_0001400 unlike FUS/QKI, which are known to regulate wide range of circRNAs (Kristensen et al., 2019), these proteins may be critical to reveal how KSHV induces specific circRNAs; the specificity of those RBPs that regulates the ratio of forward- vs back-splicing determines which circRNAs to be induced or decreased upon infection. Functional analysis of these splice factors may lead to deeper understanding of viral regulation of circRNAs, which can potentially be clinical targets.

In conclusion, we identified the circRNA that is induced by KSHV infection and promotes the growth of latently infected cells while suppressing lytic cycle early after infection. Since maintaining latent infection is crucial for oncogenic viruses, hsa_circ_0001400 may have wider implications in other viruses. Circ_0001400 is one of many host circRNAs induced upon infection and the investigation of more circRNAs will uncover this new layer of virus-host interactions via non-coding RNAs.

## MATERIALS AND METHODS

### Cell culture

Human umbilical vein endothelial cells (HUVECs) and Human Dermal Lymphatic Endothelial Cells (LEC) were obtained from Lonza and PromoCell, respectively, and passaged in EGM-2 medium (Lonza) for up to 5 passages, with passages 3 to 5 used for experiments. Media was replaced every two days. HEK-293 and 293T cells (ATCC) were maintained in DMEM medium supplemented with 10% FBS (Gibco), penicillin (100 U/ml; Gibco), and streptomycin (100 mg/ml; Gibco). The infected iSLK cell line (WT BAC16 strain) is a gift from Rolf Renne (1). They were maintained in DMEM supplemented with 10% FBS, penicillin, streptomycin, hygromycin (1200 µg/ml), puromycin (1 µg/ml), and G418 (250 µg/ml). EBV-positive and negative Akata cell lines were maintained in RPMI medium supplemented with 10% FBS, penicillin, and streptomycin. Cells were cultivated at 37 °C in a 5 % CO_2_ incubator.

### KSHV production (iSLK) and infection

KSHV BAC16 virus stock was prepared by inducing lytic cycle in iSLK-BAC16 cells with Doxycycline (1 µg/ml; Sigma-Aldrich) and sodium butyrate (1 mM; Cayman Chemical) for three days. Collected supernatants were cleared of debris with centrifugation at 500×g for 5 minutes, filtered (Rapid-Flow 0.45 µm filter; Nalgene), and pelleted with centrifugation at 30,000g for 16 hours. The virus pellet was resuspended to EBM2 (Lonza) media. Titration method was adapted from a previous report (Tagawa et al., 2016). HUVECs or LECs were infected for 3 days and the GFP positive population was quantified by CytoFLEX S (Beckman Coulter). Here, one unit of multiplicity of infection (MOI) was defined as the amount of viruses enough to infect all cells if virions are distributed equally in the culture.

*De novo* infections in HUVECs and LECs were carried out by diluting concentrated supernatant in EGM2 medium at a MOI of 1.0 unless otherwise mentioned. Polybrene (Millipore) was added at 8 μg/ml. Virus supernatant was washed off after overnight incubation and replaced with fresh EGM2. Cells were refed every 2 days until harvest.

### Microarray

Akata cells were maintained less than 1×10^6^ cells/ml. Total RNAs were extracted at log phase of cell growth at three different days. RNAs were extracted with Direct-zol RNA miniprep kit (Zymo Research) with an on-column DNase I digestion following manufacturer’s instruction. Total RNAs were subjected to the Human Circular RNA Array (ArrayStar). Total RNA from each sample was treated with RNase R to enrich circular RNA. The enriched circular RNA was reverse transcribed with random primers containing a T7 promoter. The cRNA was linearly amplified and Cy3 labeled by in vitro T7 polymerase transcription using Arraystar Super RNA Labeling protocol (Arraystar). The labeled cRNAs were hybridized onto the Arraystar Human circRNA Arrays V2 (8×15K, Arraystar), and incubated for 17 hours at 65°C in an Agilent Hybridization Oven. The microarray slides were scanned with an Agilent Scanner G2505C.

Agilent Feature Extraction software (version 11.0.1.1) was used to analyze acquired array images. Quantile normalization and subsequent data processing were performed using the R software limma package. Quantitative results are available in Table S1. Raw data for Akata cells is available at GSE206824 (Gene Expression Omnibus, GEO). For HUVECs, previously performed microarray (GSE120045, GEO) was used.

### circRNA manipulation

For depletion of circ_0001400, previously reported siRNA, as described as siCirc1400-2, was used (Tagawa et al., 2018). This siRNAs target back-splice junctions of circ_0001400 and not affect the linear counterpart RNA, *RELL1*. 2×10^5^ of HUVECs or LECs were transfected with 20 nM ON-TARGETplus siRNA (Horizon), 3 ul Dharmafect I (Horizon), and 0.4 ml Opti-MemI (Gibco) in 2ml EGM2 according to manufacturer’s guidance, followed by KSHV infection in next day. For 293T cells, 2×10^6^ cells were transfected with 20 nM siRNA with 18 μl of RNAiMax (Thermo Fisher Scientific) and 0.6ml Opti-MemI in 3ml DMEM for 48 hours.

Ectopic expression of circ_0001400 in 293T cells was done with a plasmid vector as previously reported (Tagawa et al., 2018). 2 μg pcDNA3.1- hsa_circ_0001400 of plasmids was transfected to 2×10^6^ cells with 12 μl Transporter 5 (Polysciences) lipofection reagent and 1 ml Opti-MemI in 10 ml DMEM. For primary endothelial cells, lentiviral vectors were used. pcDNA3.1-has_circ_0001400 and pcDNA3.1(+) ZKSCAN1 MCS-WT Split GFP + Sense IRES (Addgene plasmid # 69909) were digested with BamHI and XhoI (NEB) and cloned into pLV-mCHerry:T2A:Bsd-CMV plasmid (VectorBuilder). VSV-G pseudotyping and packaging was performed by VectorBuilder. Titration was performed as for KSHV BAC16, but with mCherry reporter. For infection, KSHV and lentivirus are added to 1×10^6^ HUVECs or LECs simultaneously with 8 μg/ml polybrene. Media was replaced after overnight incubation and cells.

### RNA extraction and RT-qPCR

Total RNA was extracted with Direct-zol RNA miniprep kit with on-column DNAse I digestion (Zymo Research). 0.5 to 1 μg of total RNA was used for reverse-transcription with ReverTra Ace qPCR RT Master Mix (Toyobo) and qPCR was performed with Thunderbird Next SYBR qPCR Mix (Toyobo) following manufacturer’s instructions. Following primers are used: hsa_circ_0001400, 5’- ATGTCTGTTAGTGGGGCTGA-3’, 5’- TATCTGCTACCATCGCCTTT -3’;LANA, 5’- GTGACCTTGGCGATGACCTA-3’, 5’-CAGGAGATGGAGAATGAGTA - 3’;RTA, 5’- CTGACGTCATGTCACCCTTG-3’, 5’-TCTCTACACGGCACACCTTG-3’;*TTI1*, 5’- CCCTCCATTCTGCCACGTTTA -3’, 5’- ACACTGCAAGAGTAGAACCTGTA -3’;*CNOT10*, 5’- CAGTCTTCGGCCATTCCTGT -3’, 5’- GCCCCATTTTCCTGCTTTGG -3’;*ACTB*, 5’- TCACCCACACTGTGCCCATCTACGA -3’, 5’- CAGCGGAACCGCTCATTGCCAATGG- 3’;*GAPDH*, 5’- TTCACACCCATGACGAACAT -3’, 5’- TTCACACCCATGACGAACAT - 3’;*RPS13*, 5’- TCGGCTTTACCCTATCGACGCAG-3’, 5’- ACGTACTTGTGCAACACCATGTGA -3’.

### RNA-Seq and data analysis

circ_0001400-manipulated and KSHV (BAC16)-infected HUVECs and LECs were harvested for total RNA. Library preparation and sequencing was performed by Genewiz. For circ_0001400-depleted HUVECs and ectopically expressed HUVECs, Illumina strand-specific RNA-seq with Poly-A selection and Illumina RNA-seq with rRNA depletion, respectively, were employed. Prepared libraries were sequenced with HiSeq 2500 (Illumina) in 2×150 bp paired-end conditions. Fastq files are available at GSE206928 (GEO). In addition, previously reported data are used: KSHV-infected LECs (PRJNA851845 (SRR20020770, SRR20020769, SRR20020761, SRR20020757, SRR20020758)), KSHV-infected HUVECs (GSE165328), HCMV-infected MRC5 (GSE155949) (Oberstein and Shenk, 2017)). Sequenced reads were mapped to GRCh38 with KSHV genome, NC009333, by STAR 2.7.6a, trimmed with trimmed with cutadapt 1.18 (Martin, 2011), mapped to GRCh38 with KSHV genome, NC009333, using STAR 2.7.6a (Dobin et al., 2012), and gene count files are created using RSEM (Zhao et al., 2011). Filtering, normalization, quality control, and differential expressed gene (DEG) analysis are performed using DESeq2 and NIDAP (NIH Integrated Data Analysis Platform). On NIDAP, after filtering of genes with low sequenced reads, read counts are normalized with quantile normalization for DEG analysis by voom (limma package). Pathway enrichment analysis was performed with Ingenuity Pathway Analysis (IPA, Qiagen). DEGs from both circ_0001400-depleted HUVECs and ectopically expressed HUVECs were determined by adj. p-value <0.05. Among them, only genes that were regulated in opposite way by ectopic expression and depletion were further chosen (Table S3 and S4) for IPA analysis. To quantify circRNAs and circRNA:linear RNA ratios, data from mock and KSHV-infected HUVECs were subjected to CIRCExplorer3 (Ma et al., 2019). This software bases quantitation to intron-spanning reads, only, such that reads from forward-splicing (only from linear RNAs) and back-splicing (only from circular RNAs) can be directly compared.

### 4SU RNA-Sequencing

iSLK-BAC16 cells were uninduced or induced for lytic cycle as described. At one or three days after induction, 10 mM 4SU (Sigma-Aldrich) was added to cell culture medium. 15 minutes post-4SU addition, cells were collected, and RNAs were extracted. 40-50 ug total RNA was biotinylated in 10 mM Tris pH 7.4, 1 mM EDTA, 0.2 mg/mL EZ-link Biotin-HPDP (Thermo Fisher Scientific). Unbound biotin was removed by performing a chloroform:isoamyl alcohol extraction using MaXtract High Density tubes (Qiagen). RNA was isopropanol precipitated and resuspended in water. Biotinylated RNA was bound 1:1 to Dynabeads My One Streptavidin T1 (Thermo Fisher Scientific) equilibrated in 10 mM Tris pH 7.5, 1 mM EDTA, 2 M NaCl. Bound beads were washed three times with 5 mM Tris pH 7.5, 1 mM EDTA, 1 M NaCl. 4SU-RNA was eluted with 100 mM DTT and isolated using the RNeasy MinElute Cleanup Kit (Qiagen). RNA was sent to the NCI CCR-Illumina Sequencing facility for library preparation and sequencing. Pulldown RNA was ribominus selected using the NEBNext rRNA Depletion Kit v2 (NEB # E7400L) and RNA-Seq libraries were generated using the NEBNext Ultra II Directional RNA Library Prep Kit (NEB # E7760L). Two biological replicates were sequenced using the Illumina NextSeq 550 platform to generate 150 bp PE reads. Fastq files are available at BioProject #PRJNA851589; SRA accession codes: SRR19792341, SRR19792340, SRR19792338, SRR19792337, SRR19792339, SRR19792336.

### KSHV production assay

2×10^5^ LECs were depleted for circ_001400 and infected with KSHV as described for three days in 2ml EGM media. Conditioned media was collected and filtered with Minisart 1.2 µm filter (Satorius). 1×10^5^ HEK-293 cells are seeded, and media was replaced with 1ml diluted conditioned media (1:1 with DMEM) and incubated for three days. KSHV-infected GFP-positive HEK-293 cells were measured by CytoFlex S (Beckman Coulter) flow cytometer.

### Cell growth (cell cycle and apoptosis)

2×10^5^ HUVECs were depleted for circ_001400 and infected with KSHV as described for three days in 2ml EGM2 media. Cell cycle was measured with Click-iT EdU Pacific Blue Flow Cytometry Assay kit (Thermo Fisher Scientific) and FxCycle PI/RNase staining solution (Thermo Fisher Scientific) following the manufacturer’s instructions. Cells were incubated with EdU for 4 hours before fixation and staining. Fluorescence was measure by CytoFlex S (Beckman Coulter) flow cytometer. For apoptosis assays, 1×10^4^ HUVECs were cultured in EGM2 or dilution with EBM2 (Lonza). Apoptotic cells were stained with ViaStain Live Caspase 3/7 kit (Nexcelom) and images were taken and anlyzed with Celigo Image Cytometer (Nexcelom).

### Immunostaining

2×10^5^ HUVECs were depleted for circ_001400 and infected with KSHV as described for three days in 2ml EGM media. Cells were washed with PBS, detached with Accutase (BioLegend), and mixed with antibodies in staining buffer (PBS, 0.5% FBS, 2mM EDTA) for 20 minutes at room temperature. Fluorescence was measure by CytoFlex S (Beckman Coulter) flow cytometer. Median fluorescence intensities (MFIs) were quantitated with FlowJo X (BD Biosciences) and MFIs for each antibody is scaled to corresponding isotype controls. Following antibodies were used: PE/Cyanine7 anti-human CD40 antibody (5C3, BioLegend); PE/Cyanine7 Mouse IgG1, κ Isotype ctrl antibody (MOPC-21, BioLegend); Alexa Fluor 647 anti-human CD54 antibody (HCD54, BioLegend); Alexa Fluor 647 Mouse IgG1, κ Isotype ctrl antibody (MOPC-21, BioLegend); PE anti-human CD275 (B7-H2, B7-RP1, ICOSL) antibody (9F.8A4, BioLegend); PE Mouse IgG1, κ Isotype ctrl antibody (MOPC-21, BioLegend).

### circRNA-pulldown

The method to pull-down circRNA-RNA complexes was adapted from Ziv et al., 2018 (Ziv et al., 2018). 4×10^6^ 293T cells were transfected with 2 µg of pcDNA3.1- hsa_circ_0001400 and 12 µl Transporter 5 for 24 hours. Cells were washed and replaced with 5mg of Psoralen (Berry) resolved in 10ml PBS. After 20 minutes incubation, cells were irradiated with UVA for 10 minutes (Stratalinker 1800) to crosslink RNAs. The extracted total RNA was mixed with 100 pmol of biotinylated DNA oligos (IDT) and incubated for 6 hours at 37°C in 1.5 ml hybridization buffer (500mM NaCl, 0.7% SDS, 33 mM Tris-Cl, pH 7, 0.7 mM EDTA, 10% formamide). Add 100 µl of MyOne Streptavidin C1 Dynabeads to RNA (Thermo Fisher Scientific) and incubate for an additional hour. Beads were washed with 2×SSC buffer + 0.5% SDS, and RNA was released by DNase I treatment (Ambion, 0.1 unit/µl) for 30 minutes at 37°C. RNA was purified with RNA Clean&Concentrator-5 kit (Zymo Research) followed by de-crosslinking with UVC exposure (2.5kJ/m^2^, Stratalinker 2400). SMART-Seq v4 Ultra Low Input Kit for Sequencing was used for full-length cDNA synthesis and amplification (Clontech), and Illumina Nextera XT library was used for sequencing library preparation. Illumina HiSeq 2500 was used to sequence samples with 2×150 Paired End configuration. Image analysis and base calling were conducted by the HiSeq Control Software (HCS). Raw sequence data was converted into fastq files and de-multiplexed using Illumina’s bcl2fastq 2.17 software. Fastq files are available at GSE206929 (GEO). Enriched transcripts were identified through DEG analysis on NIDAP platform as described earlier. For these samples, read counts were median scaled ant not normalized before DEG analysis. All enriched transcripts are available in Table S4. Used oligos are as follows: control (sense strand of circ_0001400), 5’- TGGCACAGAGTAGCAGCGAATGCTGATGT -Biotin-TEG-3’; probe (anti sense of circ_0001400), 5’- ACATCAGCATTCGCTGCTACTCTGTGCCA -Biotin-TEG-3’.

### CARPID of circ_0001400

CRISPR-assisted RNA–protein interaction detection method (CARPID) was performed as described by Yi et al. (Yi et al., 2020). 4×10^6^ 293T cells were transfected with 2 µg of pcDNA3.1- hsa_circ_0001400, 3 µg of CARPID BASU-dCasRx (Yi et al., 2020), 3 µg of gRNA-expressing plasmids based on pXR004: CasRx pre-gRNA (Konermann et al., 2018), and 48 µl Transporter 5 for 24 hours. CARPID BASU-dCasRx was a gift from Jian Yan & Liang Zhang (Addgene plasmid # 153209 ; http://n2t.net/addgene:153209 ; RRID:Addgene_153209). pXR004: CasRx pre-gRNA cloning backbone was a gift from Patrick Hsu (Addgene plasmid # 109054 ; http://n2t.net/addgene:109054 ; RRID:Addgene_109054). Media was replaced with fresh DMEM and incubate 24 hours more. Media was changed with DMEM + 200 µM biotin (Sigma-Aldrich) and cells were incubated for 15 minutes at 37°C. Cells were washed with 10ml three time and scraped into 1 ml ice-cold PBS. Cells were pelleted and lysed in 1mL of 0.5% NP40 lysis buffer (150 mM NaCl, 50 mM Tris pH ∼7, 0.5% NP40, and protease inhibitors) by gently pipetting the solution until fully homogenous. Cell lysates were kept on ice for 30 min followed by pulse sonication to disintegrate the DNA. Lysates were clarified using centrifugation for 10 minutes at 14000 RPM at 4°C. The supernatant was removed and incubated with 30 ul of slurry Pierce Neutravidin Agarose beads (Thermo Fisher Scientific Catalog #29200) overnight at 4°C with rotation. The following day, the beads were washed three times with ice-cold 1x-TBS before resuspending with 25mM ammonium bicarbonate (pH 8.0). The beads were placed on a heat plate for approximately 4 minutes at 95 °C and allowed to cool. Once cooled the beads were digested overnight with 2 µg of Trypsin (Promega Catalog # V5111) at 37 °C and constant rocking.

For clean up and mass spectrometry, the peptides were desalted and eluted using 70% ACN/0.1%TFA, following the procedure for the Pierce C18 spin columns (Thermo Fisher Catalog # 89873) and dried down. The dried eluates were resuspended in 0.1% TFA. Peptides were analyzed on a Q Exactive HF (Thermo Fisher Scientific) mass spectrometer coupled to Easy nLC 1000 system (Thermo Fisher Scientific) fitted with Acclaim PepMap 100 C18 LC column (Thermo Fisher Scientific). The peptides were eluted with a 5% to 36% gradient of Acetonitrile with 0.1% Formic acid over 56 minutes with a flow rate of 300 nl/min. The QE HF was operated with each MS1 scan in the orbitrap at 60,000 resolution with a maximum injection time of 120 ms and an AGC target of 1e6. The MS2 scans had a normalized collision energy of 27 and were run at 15,000 resolution with a maximum injection time of 50 ms and an AGC target of 2e5.

For data analysis, acquired MS/MS spectra were searched against a human uniprot protein database in Proteome Discoverer 3.2 (Thermo Fisher Scientific) using Label free Quantification with Minora feature detection. Significantly enriched proteins were identified with Bioconductor RankProd package(Carratore et al., 2017). Scaled signals for each sample were separated into two groups: enriching gRNAs (gCirc1400_1, gCirc1400_2, gCirc1400_3) and controls (pre-gRNA, negative control gRNA), and rank-product analysis was performed as two-class, unpaired case. Proteins with p-values < 0.05 were classified as enriched. All candidates are listed in Table S2. For specific targeting of circ_0001400, we cloned guide RNAs (gRNAs) specific to the circRNAs into CasRx pre-gRNA plasmid. gRNA sequences for circ_0001400 were adapted from Lit et al., 2021 (Li et al., 2021). CasRx pre-gRNA was digested with BbsI and oligo DNAs harboring gRNAs and BbsI-compatible overhands were ligated. To test specificities and efficacies, 1×10^5^ 293T cells were transfecting with 300 ng of cloned gRNA-expression vectors and 300 ng of pXR001-EF1a-CasRx-2A-EGFP for two days. RT-qPCR was performed to confirm the knockdown of circ_0001400 as described. pXR001: EF1a-CasRx-2A-EGFP was a gift from Patrick Hsu (Addgene plasmid # 109049 ; http://n2t.net/addgene:109049 ; RRID:Addgene_109049). Following oligos were used for cloning: negative control, 5’- AAACTCTGTAGTCGTAAGCCTGCTACTCTGTGCC -3’, 5’- CTTGGGCACAGAGTAGCAGGCTTACGACTACAGA -3’; gCirc1400_1, 5’- AAACAGACATCAGCATTCGCTGCTACTCTGTGCC -3’, 5’- CTTGGGCACAGAGTAGCAGCGAATGCTGATGTCT -3’; gCirc1400_2, 5’- AAACTCAGCATTCGCTGCTACTCTGTGCCACTGC -3’, 5’- CTTGGCAGTGGCACAGAGTAGCAGCGAATGCTGA -3’; gCirc1400_3, 5’- AAACCTTTAAGACATCAGCATTCGCTGCTACTCT -3’, 5’- CTTGAGAGTAGCAGCGAATGCTGATGTCTTAAAG -3’.

### RNA-IP of circ_0001400

12×10^6^ 293T cells were transfected with 8 µg of pcDNA3.1- hsa_circ_0001400 and 48 µl Transporter 5 for 24 hours. Cells were detached and 10 million cells was used per antibody. Cells were washed with PBS twice, lysed in 500 µl IP buffer (20 mM Tris-CL, pH 7.5, 150 mM NaCl, 1 mM EDTA, 0.5% NP-40, protease inhibitor), incubated for 10 minutes at 4°C, and pelleted. Supernatant was collected pre-cleared with 20 µl Protein A/G MagBeads (GenScript) per 1.5 ml lysate for 30 minutes, 4°C. Beads were removed and supernatant was incubated with 4 µg antibody/400 µl lysate overnight at 4°C. 70 µl magbeads were added and beads were incubated for 4 hours at 4°C. Beads were washed three times with 1ml PBS and RNAs were eluted with Trizol and extracted with Direct-zol RNA miniprep kit (Zymo Research) followed by RT-qPCR. Following antibodies were used: □-SFRS18/SRrp130 (A301-609A, Thermo Fisher Scientific); □-eIF3k (PA5-98862); Rabbit IgG Control (GenScript).

## Data analysis and availability

Most graphs contain plots with each data point represented and also include the mean and standard deviations (SD). For testing of significance, paired-t tests were performed with Prism (GraphPad) except for high-throughput analysis like RNA-Seq and microarray, and presented with asterisks indicating P values. Sequencing and microarray data produced in this work are available at Gene Expression Omnibus (GSE206930) and Sequence Read Archive (PRJNA851845, PRJNA851589).

## ACKNOWLEDGEMENTS

This work was supported by the Intramural Research Program of the Center for Cancer Research, NCI, NIH (1ZIABC011176) and Japan Society for the Promotion of Science (Research Fellowship for Japanese Biomedical and Behavioral Researchers at NIH). The funders had no role in study design, data collection and analysis, decision to publish, or preparation of the manuscript. We thank the Center for Cancer Research Sequencing Facility (Frederick, MD) for their assistance in sequencing libraries. The resources of the NIH High-Performance Computing Biowulf Cluster were utilized for all our computational needs. Figures are partially created with BioRender.com.

## LEGENDS

**Fig. S1.**
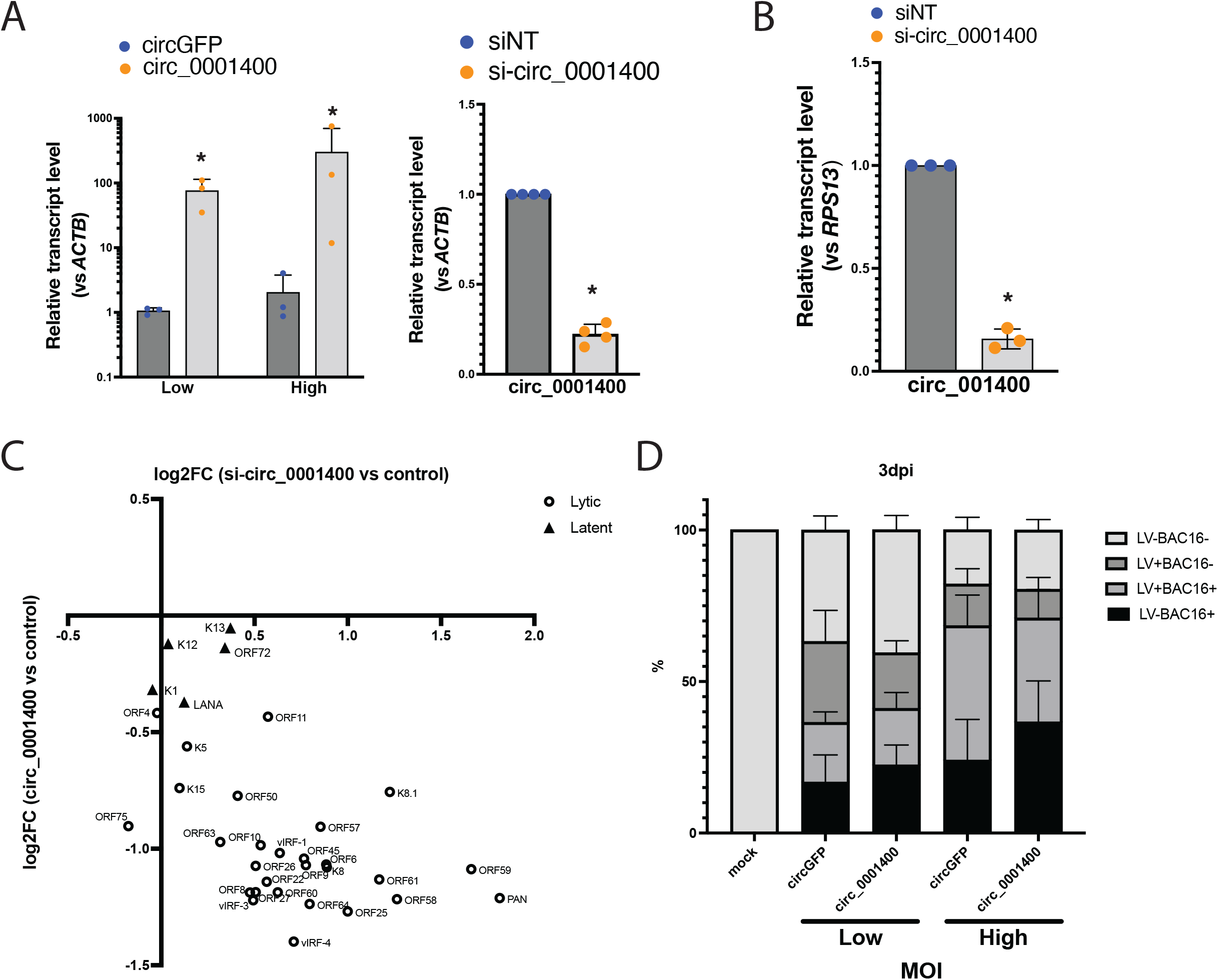
circ_0001400 affects mainly viral lytic gene expression. (A) circ_0001400 transcript levels after manipulation of circ_0001400. circ_0001400 was ectopically expressed with lentivirus (left) or treated with siRNA (right) in KSHV-infected HUVECs. MOIs were 0.5 (Low) or 1.0 (High) for lentivirus and 0.5 for the siRNA condition. Total RNAs were extracted followed by RT-qPCR at 3 days post infection (n=3). (B) circ_0001400 transcript levels after manipulation of circ_0001400. circ_0001400 was depleted by siRNAs in KSHV-infected LECs. Total RNAs were extracted followed by RT-qPCR at 3 days post infection (n=3). (C) Log_2_ fold changes of KSHV transcript levels after manipulation of circ_0001400 in HUVECs. circ_0001400 was ectopically expressed with lentivirus or depleted by siRNAs in KSHV-infected HUVECs. At 3 days post infection, total RNAs were extracted for RNA-Seq. x-axis shows the effect of circ_0001400 depletion while y-axis describes the result of circ_0001400 ectopic expression. The list of fold changes is available in Table S2. (D) Flow cytometry analysis of HUVECs dually infected with KSHV and lentiviruses. KSHV BAC16 contains a GFP reporter gene whereas circRNA-expressing lentivirus contains mCherry reporter gene. Both markers are under constitutive promoters and useful to track infection (n=3). Infectivity of HUVECs was measured at 3 days post infection with KSHV BAC16.

**Table S1. circRNA expression changes upon herpesvirus infections and circRNA-linearRNA ratios in KSHV-infected HUVECs**.

**Table S2. All candidates of circ_0001400-interacting proteins**.

**Table S3. KSHV transcriptome of circ_0001400-manipulated KSHV-infected HUVECs**.

**Table S4. Differentially expressed genes by circ_0001400 in KSHV-infected HUVECs**.

**Table S5. All candidates of circ_0001400-interacting transcripts**.

